# An Extended Clade Framework for Annotated Trees in the Context of Phylogeography and Transmission Tree Inference

**DOI:** 10.64898/2026.04.23.720428

**Authors:** Lars Berling, Caroline Colijn

## Abstract

Bayesian phylogenetic inference commonly produces samples of thousands of trees representing a posterior distribution and associated variables. Such a collection of trees is hard to interpret and it is common practice to summarize such samples into a single representative tree. Methods for constructing representative trees have largely been restricted to plain tree topologies, encoding only relationships among taxa. Inference with more sophisticated models produces annotated tree objects. These have additional information representing nodes’ locations in the case of phylogeography, host information when inferring transmission trees, or sampled ancestor status when incorporating fossil information. Nevertheless, these annotated representations are reduced to a single representative tree, typically using methods developed for plain tree topologies and without accounting for the resulting methodological mismatch. Here, we introduce the concept of an extended clade and investigate an extension of the conditional clade distribution (CCD) model. Through motivating examples and case studies in discrete trait phylogeography and transmission tree reconstruction, we demonstrate limitations of standard summary tree approaches and show how these can be addressed using an extended CCD framework that explicitly incorporates the annotated tree structure.

## Introduction

Bayesian phylogenetic inference using Markov chain Monte Carlo (MCMC) methods produces large samples from a posterior distribution over phylogenetic trees. These samples represent uncertainty in both tree topology and associated parameters, and typically consist of thousands of trees, making direct interpretation difficult. As a result, it is common practice to summarize such samples using a single representative tree for visualization, interpretation, or downstream analysis.

A large body of work has investigated how to construct a representative tree from collections of trees, leading to classical methods such as the majority rule consensus (MRC) tree (Margush and McMorris, 1981) and the maximum clade credibility (MCC) tree; see Bryant (2003) for an overview. More recently, this has been extended by methods such as the highest independent posterior subtree reconstruction (HIPSTR) (Baele et al., 2025a) and the conditional clade distribution (CCD) MAP tree (Berling et al., 2025). The MRC tree has also recently been revisited by Takazawa et al. (2026), who propose fine-grained dissimilarity measures for clades that outperform the classical approach. However, these approaches are almost exclusively designed for plain tree topologies, that is, trees encoding only relationships among taxa. In such settings, summarizing is typically performed in two steps: first selecting a sampled topology or estimating a representative topology, and then annotating it with marginal summaries such as branch lengths or posterior support.

In many applications, however, phylogenetic inference is performed jointly with additional model components, resulting in annotated trees that carry richer structure. Examples include phylogeography (Vaughan et al., 2014; Kühnert et al., 2016; Vaughan and Stadler, 2025), where nodes are associated with geographic locations; phylodynamics and epidemiology, where trees also encode transmission or population processes (Didelot et al., 2017; Duault et al., 2022; Colijn et al., 2026; Specht et al., 2025); fossil-calibrated inference with sampled ancestors (Gavryushkina et al., 2014; Stolz et al., 2025); and models allowing for reticulation or hybridization (Zhang et al., 2018; Müller et al., 2020; Neureiter et al., 2022). These joint models aim to propagate uncertainty coherently across interacting variables, capturing dependencies between the phylogeny and associated implied structure.

Despite this, annotated posterior tree samples are still commonly summarized using methods developed for plain tree topologies, followed by ad hoc overlay of annotations. This decoupled summary procedure ignores the joint structure of the inference problem and can lead to summaries that are internally inconsistent, misleading, or incompatible with the underlying model.

In this paper, we address this conceptual gap by introducing the notion of an extended clade, providing a general framework for summarizing posterior distributions over annotated trees. We primarily investigate an extension of the conditional clade distribution (CCD) model (Höhna and Drummond, 2012; Larget, 2013; Lewis et al., 2016; Berling et al., 2025). Building on that, we show how extended clades can also be used to define generalized versions of classical tools such as the Robinson-Foulds (RF) metric and the majority rule consensus tree. Through case studies in discrete trait phylogeography and transmission tree inference, we illustrate how standard summary tree approaches can fail in the presence of annotated structure, and demonstrate how the extended CCD framework enables coherent and model-consistent summaries.

Branch lengths could in principle also be treated as annotations within this framework. However, unlike the discrete annotations we consider, branch lengths are continuous, and discretising them would require additional assumptions we prefer to avoid. We therefore focus on annotations that are already discrete, and leave the incorporation of continuous annotations such as branch lengths to future work.

### Extended Clades and CCDs

We first recall definitions of previous work (Berling et al., 2025) and then define our extended clade framework. Throughout this section we consider phylogenetic trees on the set of taxa *X*.

### Clades

The internal nodes of a rooted tree on the set of taxa *X* form a hierarchy. Formally, a hierarchy *H* on the set *X* is a collection of non-empty subsets of *X* with the property that any two elements of *H* are either nested (one is contained in the other) or disjoint (Steel, 2016). Given a rooted phylogenetic tree *T*, and a vertex *v* of *T*, the clade associated with *v*, denoted *c*_*T*_ (*v*), is the subset of *X* that is descendant from *v*. We will let

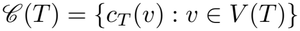

denote the set of clades associated with the vertices of *T*. This collection *C* forms a directed graph and it encodes the rooted phylogenetic tree *T*. In the following, we mostly drop the notation *c*_*T*_ (*v*) and refer to a clade simply as *C*.

### Forest network

A forest network *N* on a set of taxa *X* is a rooted bipartite directed graph with vertex set *V* = (*C, S*) that satisfies the following properties:

- Each node *C ∈ C* represents a clade on *X*.
- Each node *S∈ S* represents a clade split, connecting a parent clade to its two child clades via a clade split. Specifically, each split *S∈ S* has degree three with one incoming edge (*C, S*) from the parent clade and two outgoing edges (*S, C*_1_), (*S, C*_2_) to its child clades, such that *C*_1_ *∪C*_*2*_ = *C, C*_*1*_ *∩C*_*2*_ =*∅* for some *C*_1_, *C*_2_, *C∈ C*. We say that *S* = {*C*_1_, *C*_2_} is a clade split of *C*. We also use the notation *C*_1_||*C*_2_ for *S*.
- Each non-leaf clade has outdegree at least one and each clade except *X* has indegree at least one.

In this case *C*_*ρ*_ = *X* is the root of *N* and each non-leaf clade has at least one clade split. Note that, for a clade split {*C*_1_, *C*_2_} of *X*, the network *N* contains all trees composed (amalgamated) of one subtree from *N* (*C*_1_) and one subtree from *N* (*C*_2_); this holds recursively. Hence, a forest network is suitable to represent a large collection of trees when all combinations of subtrees are included.

### CCD graph

In order to turn a forest network into a tree distribution, we need to be able to compute a probability for a tree *T*. Larget (2013) suggested to use the product of clade split probabilities over all clade splits in *S* (*T*) as the probability of *T*. We define a CCD graph as a forest network *G* where each clade split *S* in *S* (*G*) has an assigned probability Pr(*S*) such that, for each clade *C* ∈ *C* (*G*), we have Σ _*S*∈*C*(*C*)_ Pr(*S*) = 1. So for a tree *T* displayed by *G*, we have

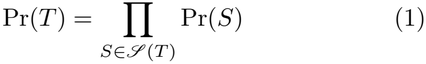

and, for any other tree *T*′, we have Pr(*T*′) = 0. Furthermore, the sum of probabilities of all trees displayed by *G* is one, see Larget (2013, Appendix 2). The resulting CCD graph is what we call a CCD1 (Berling et al., 2025).

### CCD-based point estimators

For a CCD1, we call the tree *T* with maximum probability Pr(*T*) in the CCD1 the CCD1-MAP tree. Using the recursive relationships for CCDs, we can find the CCD1-MAP tree efficiently as follows. Let Pr^*⋆*^(*C*) denote the maximum probability of any subtree rooted at clade *C*. With Pr^*⋆*^(*l*) = 1 for a leaf *l*, we can compute Pr^*⋆*^(*C*) with the following formula:

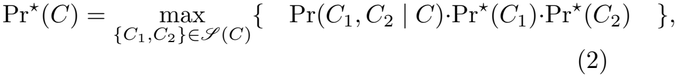

where Pr(*C, C*_1_| *C*_2_) denotes the probability of the clade split *S* = *C*_1_ ||*C*_2_. The maximum probability of any tree in the CCD1 is then given by Pr^*⋆*^(*X*). The tree *T* achieving this maximum probability can be obtained along with the corresponding value by dynamic programming.

### Extended Clades

Let *A* be a set of annotations, where each annotation *a∈ A* is associated with a node *v* in an annotated tree *T*. An extended clade is then a pair (*C, a*), where *C* = *c*_*T*_ (*v*) for some vertex *v∈ V* (*T*) and *a∈ A*. In contrast to standard clades, which capture only tree topology, extended clades incorporate additional information attached to the corresponding node or edge. We denote the set of extended clades of an annotated tree *T* by

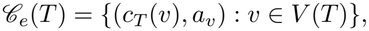

where *a*_*v*_ *∈A* is the annotation associated with vertex *v*.

This definition is intentionally general and admits different instantiations depending on the application. In discrete trait phylogeography, *a*_*v*_ corresponds to the inferred state (e.g. geographic location) of the clade *c*_*T*_ (*v*), and the set *A* is the set of discrete states in the model. In transmission tree inference, *a*_*v*_ encodes the identity of the infector that is present at the time of the node, embedding the transmission tree structure within the extended clade representation. Figure 1 illustrates both settings and their corresponding sets of extended clades.

**Fig. 1:**
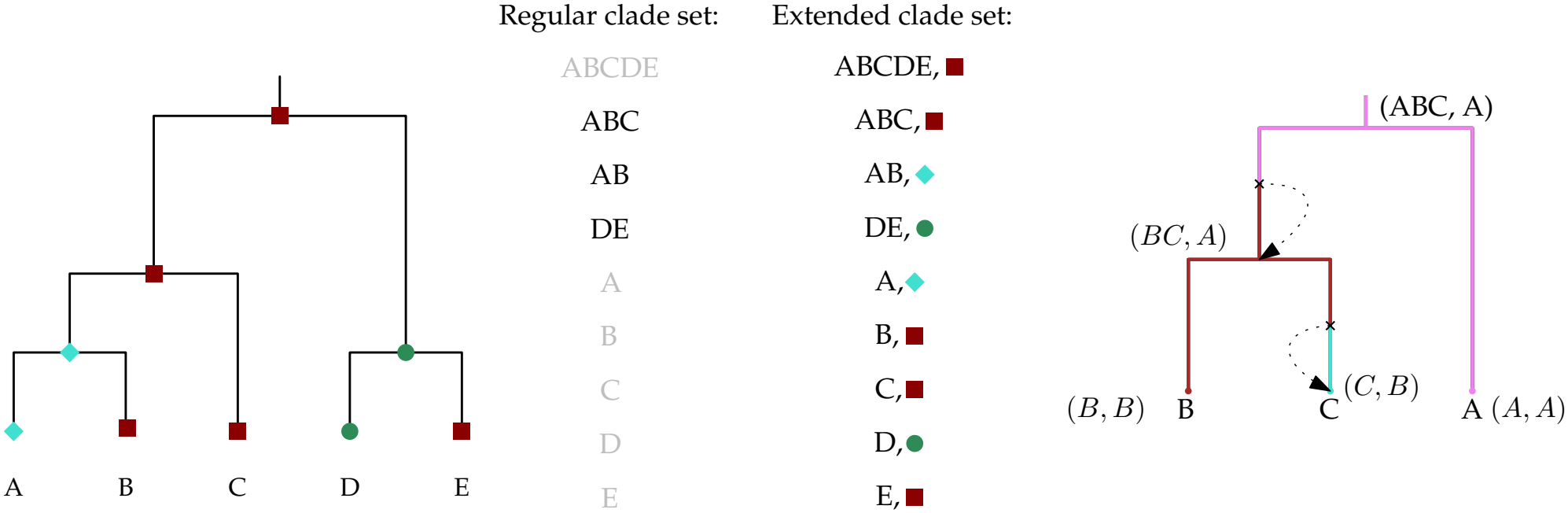
On the left, an example phylogeography annotated tree with regular and extended clade set. On the right, a transmission embedded tree with its extended clades. In both cases, we write clades as strings of leaf labels for brevity, e.g. *ABC* denotes the clade *{A, B, C}*.

In the case of discrete trait phylogenetic inference the set *A* is equal to the set of discrete states in the model. The inferred trees already contain this state information as node properties which makes the extraction of the extended clades straightforward. In the case of the transmission embedded phylogenies from the BREATH (Colijn et al., 2026) model, we get *A* = *X* ∪ {Unknown}. Each transmission represented in the tree is either between sampled individuals or unsampled hosts. Note that we do not distinguish different unsampled hosts. In this case, the extraction of the extended clades requires additional processing to reconstruct the transmission tree that is embedded in the annotated phylogenetic tree.

### Extending Conditional Clade Distributions

Given a collection of annotated trees *T* = *{T* _1_, *T* _2_, …, *T*_*k*_ *}* from a posterior sample, the CCD framework extends naturally by replacing standard clades with extended clades. Let *C*_*e*_ and *S*_*e*_ denote the sets of extended clades and extended clade splits observed in *T*, where an extended clade split of (*C, a*) is a pair {(*C*_*1*_, *a*_*1*_), (*C*_*2*_, *a*_*2*_)} such that *C*_*1*_ ∪ *C*_*2*_ = *C* and *C*_*1*_ *∩C*_*2*_ = *∅*. The forest network induced by *T* then has vertex set *C*_*e*_ ∪ *S*_*e*_ with edges naturally induced by the extended clade splits. If a tree *T* has the extended clades (*C, a*), (*C*_1_, *a*_1_), (*C*_2_, *a*_2_), and the corresponding extended clade split *S* = (*C*_*1*_, *a*_*1*_) || (*C*_*2*_, *a*_*2*_), then the forest network has the edges ((*C, a*), *S*), (*S*, (*C*1, *a*_1_)), and (*S*, (*C*_2_, *a*_2_)). Since extended clades carry annotations, the root clade *X* may appear with different annotations across the posterior sample, yielding a set of extended root clades {(*X, a*) : *a∈ A}* observed in *T*. To retain the forest network structure, namely a rooted bipartite graph, we introduce a root node *ρ* with outgoing edges to each observed extended root clade (*X, a*).

For an extended clade (*C, a*) *∈ C*_*e*_ and an extended clade split *S ∈ C*_*e*_(*C, a*), let *f (*(*C, a*)) and *f* (*S*) denote their respective frequencies in *T*. The conditional clade probability of *S* is then defined as Pr(*S*) = *f* (*S*)*/f* ((*C, a*)). For each observed root clade we set Pr(*X, a*) = *f (*(*X, a*)) */k*, where *k* = |*T* | is the total number of trees in the sample. Note that Σ_*a*_*Pr*((*X,a*)) =1 by construction. Leaf clades are not treated analogously because their annotations are handled implicitly by the hierarchical structure of the forest network and introducing these leaf probabilities would result in double-counting the normalization, as leaf frequencies are already implicitly accounted for through the clade split probabilities.

The resulting structure is what we call an extended CCD1. Note that for each (*C, a*) *∈ C*_*e*_

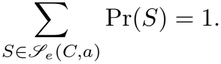

For an annotated tree *T* displayed in the extended forest network *G*, analogous to Equation 1, we have

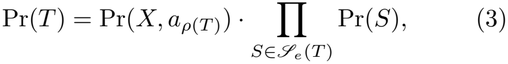

where *a*_*ρ*(*T*)_ refers to the root annotation of the tree *T*. Because the framework relies exclusively on extended clades and extended clade splits observed in the posterior sample, every tree displayed by the extended forest network is guaranteed to consist only of jointly supported topology–annotation combinations, preventing the representation of annotated trees that are incompatible with the joint posterior structure.

### CCD0 and Expansion

So far, we have focused on the extended CCD1 model, which is constructed directly from observed extended clade and clade split frequencies. In (Berling et al., 2025), we introduced the alternative model called CCD0 which differs from the CCD1 model in two ways. Firstly, its probabilities are derived from observed clade frequencies and are independent of clade split frequencies. Secondly, it features an expansion step of the underlying CCD graph. In this step, compatible clades are combined to form new clade splits which were not explicitly observed in the input tree sample.

This second step is optional but interpolates many possible splits without them being sampled which is a contributor to the good performance of the point estimator on very small sample sizes. To expand two observed clades into a new unobserved clade split a simple set comparison defines compatibility. Two classical clades *C*_*1*_ and *C*_*2*_ are considered compatible if *C*_*1*_*∩ C*_*2*_ = *∅*and *C*_*1*_ ∪ *C*_*2*_ = *C ∈ C*.

However, set-based compatibility is insufficient for extended clades as the annotation carries additional information that needs to be accounted for. For two extended clades (*C*_1_, *a*_1_) and (*C*_2_, *a*_2_) the expansion step would introduce the extended clade split (*C*_1_, *a*_1_)||(*C*_2_, *a*_2_) and connect it to all observed extended clades (*C*_*1*_ ∪ *C*_*2*_, *a*). Therefore, for each observed annotation *a* of the clade *C*_*1*_ ∪ *C*_*2*_ this introduces up to two possible new transitions, namely from *a* to *a*_1_ and *a* to *a*_2_. These possibly newly introduced annotation transitions may contradict model constraints, for example when a transition from *a* to either *a*_1_ or *a*_2_ is impossible. We illustrate the expansion for regular and annotated topologies in Figure 2.

**Fig. 2:**
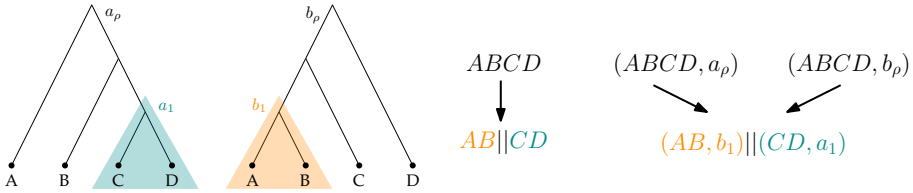
Example of an expansion step for two observed trees on the left. The regular topology extension will introduce one clade split below the root clade, displayed in the middle. For the extended clades and extended CCD on the right we have two different extended root clades with different annotations *a*_*ρ*_ and *b*_*ρ*_ where both will now be connected to the new extended clade split.

In the following paragraphs we will discuss the expansion step for the two examples we showcase in this paper. We end with a short discussion on the additional computational complexity of the CCD0 approach using extended clades.

### Phylogeography expansion

When expanding an extended CCD graph for a set of extended clades that follow a discrete state transition model from phylogeographic inference, there is no general set of rules for when two extended clades are compatible or not. The most relaxed version of such a rule, with regard to the state transition model, would be to simply use the same approach as for the regular CCD0 disregarding the clade annotations. However, with this approach we might lose the property of only supporting allowed state transitions.

Hence, a more appropriate approach might be to define compatibility with the underlying transition model in mind. This means two clades (*C*_1_, *a*_1_) and (*C*_2_, *a*_2_) are compatible if there is at least one clade (*C*_*1*_ ∪ *C*_*2*_, *a*) for which the transitions *a → a* and *a → a* are possible. We can also constrain when a transition *a→ b* is considered possible. The most relaxed way is to say we can reach state *b* from state *a* or more constraint would be to say *b* is reachable from *a* in *k* transitions, implying restriction on the number of state changes along edges in the trees.

In Figure 3 we have illustrated the possible problems with newly introduced state transitions in the expanded CCD graph and its dependency on the underlying state transition model.

**Fig. 3:**
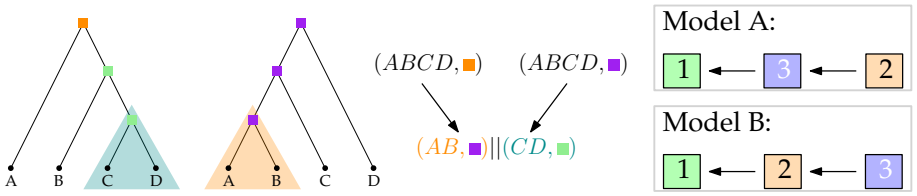
Example of an expansion step for two trees annotated with discrete states. Both sampled trees are compatible with either Model A or B. Connecting the root clade with state 3 (purple) to the new clade split is possible under either model. However, the root clade with state annotation 2 (orange) would imply a new state transition from state 2 (orange) to state 3 (purple) which is only valid in Model A and not in Model B.

In general, the CCD0 expansion step for phylogeography has to be dependent on the underlying state transition model to keep its property of only displaying valid state transitions on all trees.

### Transmission annotations from BREATH

In contrast to the broad category of possible phylogeographic models, BREATH (Colijn et al., 2026) is a single, well defined model for which we can define an explicit set of constraints for creating new extended splits from valid combinations of extended clades. These mandatory conditions for valid transmission-embedded phylogenies arise from the underlying constraints of how transmission is modelled along the phylogeny.

Recall from Figure 1 that we define the annotation such that it represents the transmission event occurring along the edge above that node and we have *A* = *X* ∪ {Unknown}. If there is no transmission event above the node (e.g. the root) then the annotation denotes the individual present at that node, which might be an unknown host. Where a direct transmission occurs along the edge, the annotation instead encodes the infector of the clade below. In the case of a block of multiple unsampled individuals above a node its annotation will always be “Unknown”.

These constraints of the underlying model must be respected when expanding the extended CCD graph with new extended clade splits. In particular, transmission paths are not permitted to overlap, since at any given point on the tree only one individual can be present; an overlap would imply that multiple individuals occupy the same point on the tree. Each individual is sampled exactly once, and for an individual to infect another clade or leaf elsewhere in the tree, the path connecting its own tip to the point of infection must be consistently annotated with this specific individual. Formally, these requirements are captured with the following conditions when adding a new extended clade split (*C*_1_, *a*_1_)||(*C*_2_, *a*_2_) to the extended parent clade (*C, a*):

Throughout we assume *i, j ∈ {*1, 2*}, i≠ j*

1. If *a*_*i*_ *∈ C*_*j*_ then *a* = *a*_*j*_ = *a*_*i*_
2. If *a*_1_ *∈ X\*(*C*_1_∪*C*_2_) then *a*_2_ = *a*_1_ or *a*_2_ = Unknown
3. If *a*_*i*_ *∈ C*_*i*_ then either *a*_*j*_ = *a*_*i*_ or *a*_*j*_ = Unknown
4. If *a ∈ C*_*i*_ then *a*_*i*_ = *a*

In Figure 4 we illustrate a valid and invalid expansion on a simple two-tree example.

**Fig. 4:**
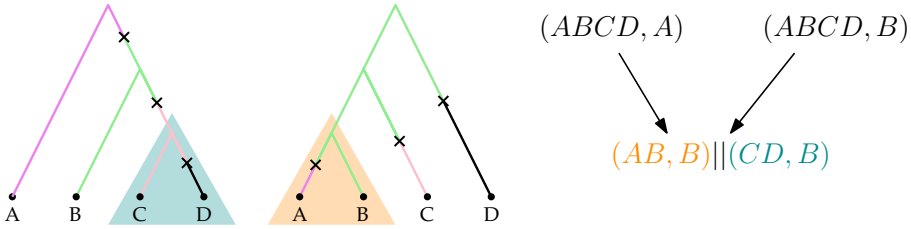
We reuse the previous two trees and clades for our example expansion step. We have two clades *C*_1_ = ({*A, B}, B*) and *C*_*2*_ = ({*C, D*}, *B*) that are combined into a new clade split. Because *B ∈C*_*1*_ we can apply rules 1 and 3 for its expansion. In addition, we can also apply rule 4 to either of the two extended clades at the root with taxa {*A, B, C, D}*. Given these rules, we can see that only one of the two possible new edges can be added to the CCD graph, namely from the extended clade (*{A, B, C, D}, B*) to the new clade split *C*_1_||*C*_2_.

### Computing CCD0 probabilities

The recursive computation of CCD0 probabilities is fixed-parameter tractable in the number of observed clades and clade splits (see Supplement of (Berling et al., 2025)). The combination of the expansion step and the recursive probability computation means that the runtime grows substantially for large trees, where the number of possible clades and new clade splits increases quickly. This is consistent with the slower CCD0 runtimes reported by Baele et al. (2025a). Substantial software optimization in the BEAST2 CCD package makes CCD0 computation feasible for such large trees.

For our extended CCDs, however, larger runtimes already arise on small trees. Since a single regular clade can now exist in multiple annotated versions, the number of possible extended clade splits multiplies accordingly, making the recursive computation of CCD0 probabilities time-consuming even for trees much smaller than those the classical CCD0 handles. This is especially apparent in the absence of the kind of software-architecture work that made the BEAST2 package efficient on large trees.

### Extended Robinson-Foulds Metric

The Robinson-Foulds (RF) (Robinson and Foulds, 1981) distance between two trees *T, T*′, denoted by *d*_*RF*_ (*T, T*′), is defined as the number of clades in one tree but not the other, and can be written as

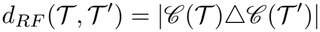

where *△* is the symmetric difference operator (i.e. *A △B* = *A ∪ B−* (*A ∩B*)). The RF metric is widely used in applications due to several advantages: it can be computed quickly (in polynomial time), its interpretation is straightforward, and its basic properties are easily established.

For two extended trees *T, T*′, the extended RobinsonFoulds (eRF) distance *d*_*eRF*_ (*T, T*′) can be defined analogously as the number of extended clades in one tree but not the other, and can be expressed as

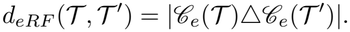

Extended clades convey information for both the tree topology and the annotated structure, hence this extension of the RF metric accounts for differences in both.

Due to these differences, the extended RF metric has slightly different properties. For example, when considering the maximum distance between any two rooted trees *T* and *T*′ with *n* leaves, the maximum RF distance max *T, T*′ *d*_*eRF*_ (*T, T*′) = 2(*n −*1)is achieved by any two trees that differ in all their non-trivial clades. The number of these clades is equal to the number of internal nodes of the tree besides the root, which is *n −*1.

For our extended clade version the maximum distance for general extensions, meaning that neither root nor leaf annotations are fixed, is given by max *T, T*′ *d*_*eRF*_ (*T, T*′) = 2(2*n −*1). This maximum distance can for example be achieved for two trees with the same topology but different annotations on all nodes including the root and leaves. If the leaf annotations are fixed, as is commonly the case in phylogeography, the maximum distance reduces to 2(*n−* 1). Interestingly, the maximum distance is independent of |*A*| but if |*A*| is large the distances are expected to be larger due to the larger number of possible mismatches.

The aforementioned example of the maximum extended RF distance immediately suggests a refinement of this metric. Instead of using the symmetric difference *△*, use a scoring function of similarity between pairs of extended clades. If two extended clades have the same classical clade but a different annotation then they contribute less to the total distance than if both classical clade and annotation are different. Furthermore, the distance could be refined using previous work on the integration of matchings to compare clades (Lin et al., 2011; Bogdanowicz and Giaro, 2011) or information theoretic extensions presented by Smith (2021). In addition, similar refinements for the annotations could also be considered. For example, annotations reflecting geographic proximity could inform the scoring, such that the distance between the same clade annotated with Germany and Canada is larger than between Germany and Belgium.

Another previous attempt at extending the RF metric to labelled trees was presented by Briand et al. (2020). However, this approach was based on tree edit operations and led to a harder problem, the computational complexity of which remains an open problem (Briand et al., 2020). Under the assumption that extended clades of an annotated tree can be computed efficiently, the extended RF metric has the same computational properties as the original version.

### Extended Clade Majority Rule Consensus

The majority rule consensus tree (MRC) contains exactly those clades that appear in more than half of the input trees. This type of tree summary is prone to polytomies, that is, parts of the tree are unresolved. A refined version of this is the greedy majority rule consensus tree, which greedily builds up a tree based on clade frequency by iteratively including clades that are compatible with the current tree and breaking frequency ties arbitrarily (Bryant, 2003). This method is occasionally referred to as the extended MRC, which we will avoid due to the possible confusion with extended clades in this paper. Despite this tree being a more resolved version of the MRC tree, the algorithm is not guaranteed to produce a fully resolved tree.

Using extended clades, we can define an analogous notion of an extended clade majority rule consensus tree. However, the extended clade setting introduces additional considerations that do not arise in the classical case. For example, root and leaf clades are classically trivial, by definition they are in every tree, but for extended clades this is not guaranteed to be the case. As a result, compatibility constraints and frequency tiebreaking may require special treatment. In particular, compatibility must be checked between the partial tree and each candidate extended clade. Despite these differences, an extended clade MRC and a greedy extended clade MRC can both be defined analogously to their classical clade counterparts.

The fine-grained dissimilarity measures recently proposed by Takazawa et al. (2026), which outperform classical MRC tree construction, could similarly prove valuable for the extended MRC tree.

### Motivating Examples

We present two examples that illustrate conceptual limitations and the methodological mismatch of standard tree summary approaches when used on annotated posterior tree distributions. For both examples, a tree topology is first selected or estimated and then in a second step annotated with summary statistics of the annotations. We show how this decoupled procedure can lead to misleading or invalid summary trees when the phylogenies are inferred jointly with their annotations.

### Discrete Trait Phylogeography

We consider a simple discrete trait phylogeographic model with three ordered states, where transitions are only allowed in one direction (i.e. the state may decrease but never increase). Locations are inferred jointly with the phylogeny under this constrained compartmental process. For illustration, we assume a posterior distribution consisting of ten sampled trees, shown in Figure 5a. A standard summary workflow, as implemented in TreeAnnotator for BEAST X (Baele et al., 2025b) and BEAST 2 (Bouckaert et al., 2019), proceeds in two steps. First, a representative topology is computed, in this case a CCD1-MAP tree. Second, the selected topology is annotated with edge and node annotations, using an attribute-specific summary statistic over all occurrences of an edge or node. Examples of these are the mean branch lengths for edges, or, for discrete state annotations, the most frequently observed state at each node. In our example, we consider only the discrete state annotations on internal nodes with the possible values {1, 2, 3}.

The outcome of this two-step procedure on our example posterior is illustrated in Figure 5b. In this example, the resulting annotated tree contains a state transition that is incompatible with the underlying compartmental model. Such a transition is absent from every tree in the posterior sample and impossible given the compartmental model, yet it appears in the annotated summary tree.

**Fig. 5:**
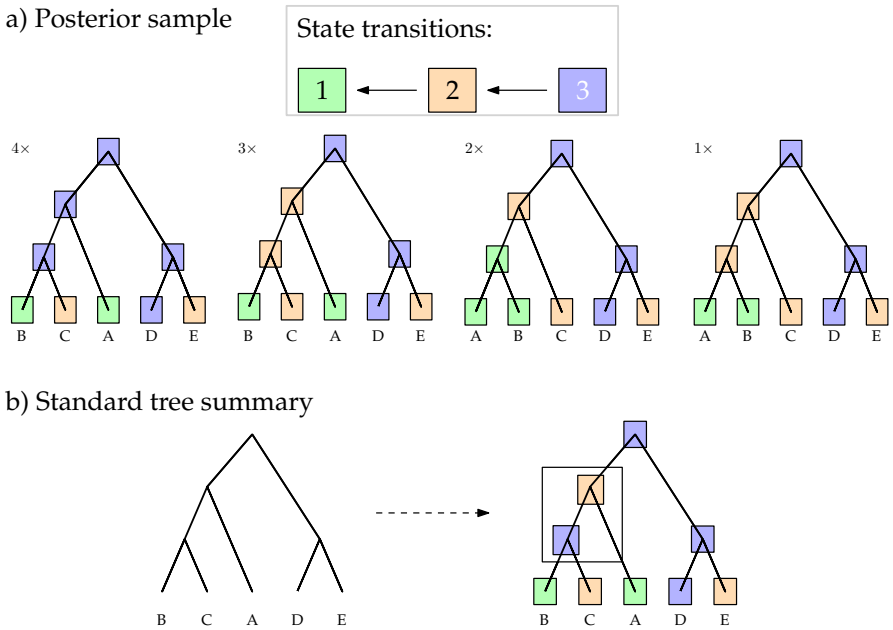
A discrete state transition model and a corresponding posterior sample. a) Example state transition model and posterior sample consisting of 10 trees with 4 different annotated topologies. Trees are weighted by their posterior frequency: 4, 3, 2, and 1 respectively. b) Result of the two-step summary applied to the posterior in panel a), highlighting an invalid state transition from state 2 to state 3. The lower panel illustrates a standard summary approach, first extracting the most likely topology using for example a CCD1 approach, then annotating most likely states on the internal nodes using the most frequent state for each node.

This issue arises because the two-step approach disregards the joint dependence between topology and node annotations. By first fixing a topology and subsequently overlaying node-specific summaries, the procedure combines quantities that were inferred jointly but summarized independently. While this strategy is consistent in settings where annotations are inferred conditional on a fixed tree, it is conceptually inconsistent when joint inference of the topology and the trait evolution is performed. In such cases, decoupled summaries can produce trees that are not supported by the posterior distribution. Furthermore, the annotated summary tree may violate structural constraints of the underlying model.

We construct an extended CCD, shown in Figure 6a, for the annotated posterior sample shown in Figure 5a. The extended CCD1-MAP tree for this example is displayed in Figure 6b and unlike the two-step procedure, it does not contain any invalid state transitions. In this example, the extended CCD does not display unsampled trees nor does it have multiple root annotations.

**Fig. 6:**
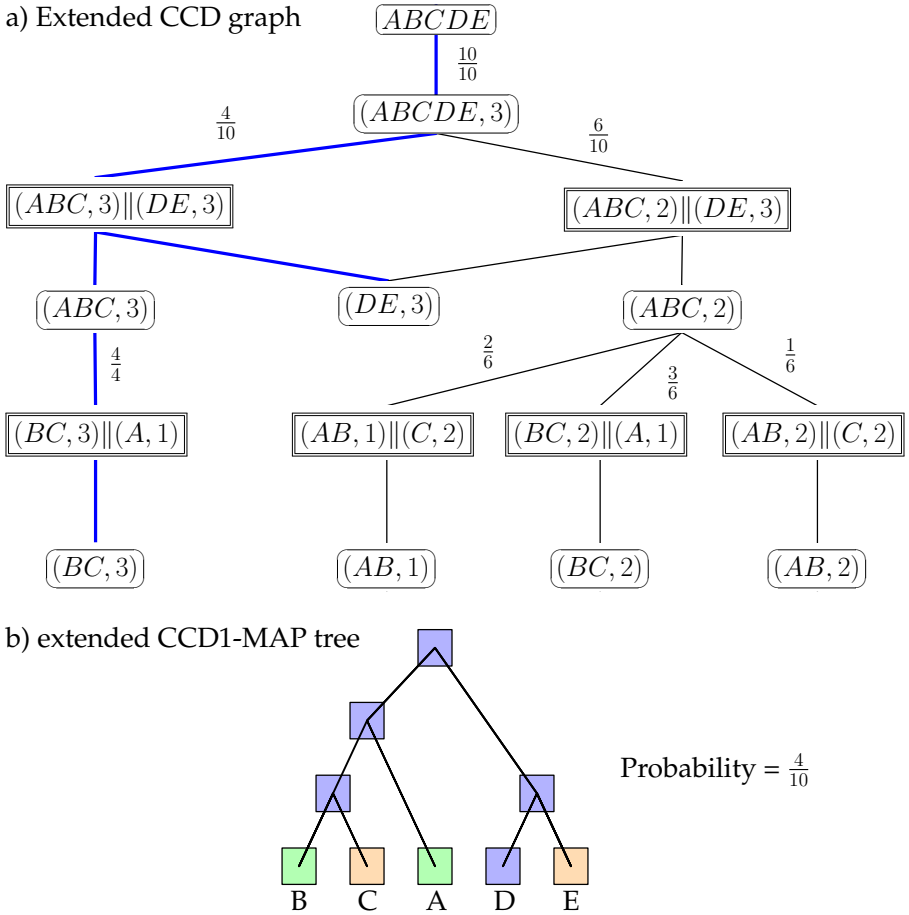
On the top, an extended CCD1 graph for the example phylogeographic posterior sample. For simplicity we have omitted cherry splits and removed set brackets from the clades. On the bottom, the corresponding extended CCD1-MAP tree, highlighted in the extended CCD1 graph on the top.

### Transmission Tree Reconstruction

In addition to the phylogeographic example, we consider phylogenies annotated with transmission trees. Here, the phylogeny inferred from sequence data is augmented with a transmission tree that specifies who infected whom. While some methods construct a phylogeny first and subsequently infer transmission events, we focus on approaches that jointly infer phylogeny and transmission structure from the data. We consider posterior samples generated by the BREATH package for BEAST 2 (Colijn et al., 2026).

Transmission events are encoded via an edge attribute blockcount and an example of such a tree is shown in Figure 7.

**Fig. 7:**
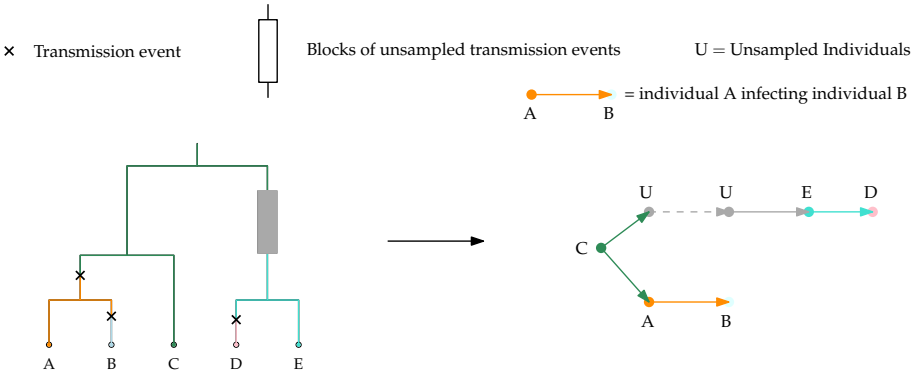
An example of a transmission embedded phylogeny (left) and the embedded transmission tree (right).

We now investigate how standard tree summary strategies perform on such posterior samples. Consider the example posterior collection shown in Figure 8a.

A naive summary strategy mirrors the common twostep approach: first compute a representative tree topology, and then overlay average summaries of the edge attributes. The resulting annotated tree is shown in Figure 8b. Because transmission events are treated as independent edge annotations, the two-step approach can result in summaries that violate structural constraints of valid transmission trees. For example, the two-step approach may fail to ensure the required presence of at least one transmission event along paths between sampled individuals (leaves). Furthermore, regular averages such as mean or median are not applicable for the discrete blockcount attribute. Summarizing these attributes requires case-specific implementations which do not currently exist.

**Fig. 8:**
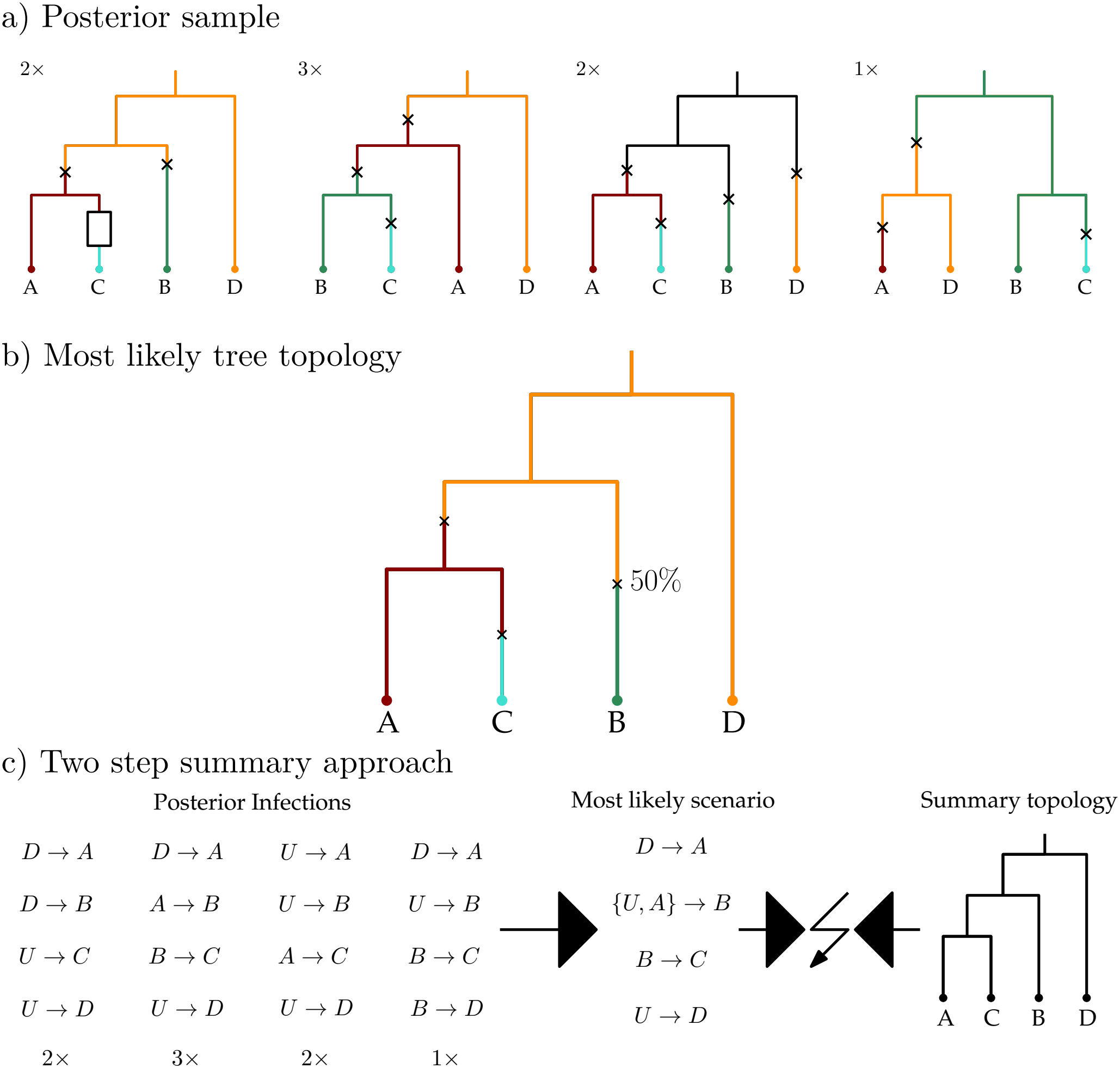
A posterior sample of transmission embedded trees: the naive summary approach and the more sophisticated but incompatible approach. a) An example posterior collection of transmission embedded phylogenies. Transmission trees are coloured in. b) Most likely tree topology annotated with average edge annotations. Picking the wrong 50% for the edge towards B (in this case no infection event) would mean that the tree becomes invalid since there would be a direct path between B and D. Or even worse, picking a mean annotation would result in an invalid blockcount value of 0.5. c) The posterior infection scenario for the example posterior in panel a). For each of the four individuals (leaves) we extract the most frequent infector from all the posterior samples. We assume that there is an unsampled infector present at the root of the tree. On the right we have the summary tree topology which is incompatible with the most frequent infection scenario. This is because the transmissions from D to A and from B to C overlap in the tree and can therefore not both be embedded.

A more refined strategy is to summarize the phylogenetic topology and the transmission trees separately using appropriate methods for each component, and then combine the two summaries. This better preserves the internal consistency of the transmission tree structure, though it still assumes independence of the two components.

The result of this separate summary is shown in Figure 8c. Here, the most likely transmission scenario extracted from the posterior is incompatible with the summarized tree topology. In particular, the transmission structure cannot be embedded into the representative phylogeny because two transmission paths are overlapping.

These examples illustrate a structural failure of existing summary approaches. Both summary strategies treat the phylogeny and the embedded transmission tree as independent objects. However, under joint inference they are tightly coupled: transmission constraints restrict the set of admissible phylogenies, and the phylogeny constrains feasible transmission histories. Decoupled summaries therefore risk producing annotated trees that are not supported by the posterior distribution and may not correspond to any valid transmission scenario.

We now construct an extended CCD1 for the posterior sample of transmission trees shown in Figure 8a. Unlike the phylogeography example, multiple extended root clades with different annotations are present in this example. Furthermore, leaf annotations are not fixed; consequently, cherry splits need to be considered and are not necessarily unique. The resulting extended CCD1 is illustrated in Figure 9a. For visual clarity, leaf clade nodes are omitted but are implied by the extended clade splits.

**Fig. 9:**
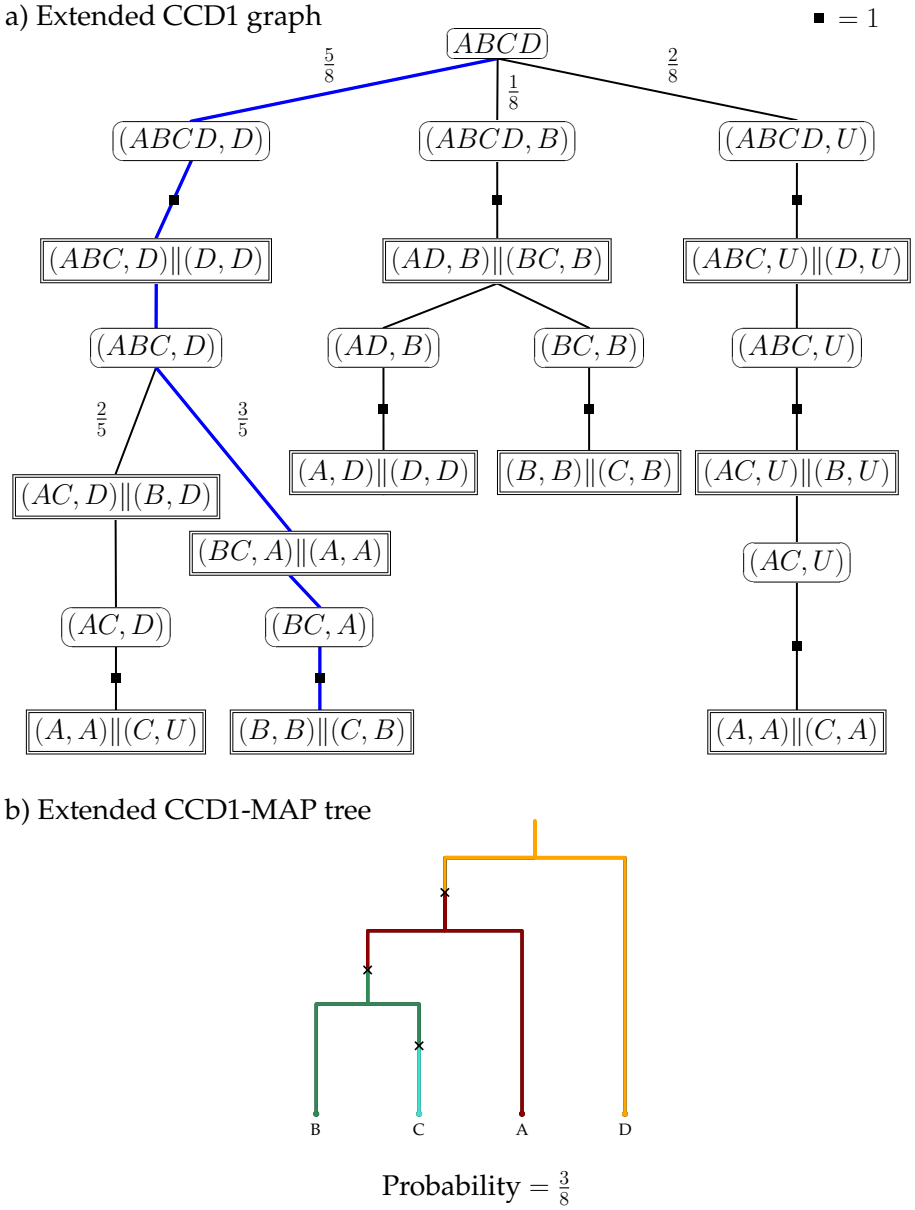
a) Extended CCD for the transmission tree posterior sample in Figure 8a. We omit the nodes corresponding to leaf clades for visual clarity; the same leaf taxon may appear with different transmission annotations. We have added squares on edges to denote extended clade splits with a conditional frequency of 1 given their parent clade. b) The corresponding extended CCD-MAP tree, its corresponding path highlighted in a).

By construction, each tree represented in the extended CCD satisfies the structural constraints of the transmission tree model, and all transmission events are consistent with the posterior sample. In this example, all trees displayed by the extended CCD correspond to the four different sampled trees, each with their sample frequency as their CCD probability. Consequently, the extended CCD-MAP tree is the second (from the left) sampled tree with a probability of 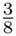.

Due to the way we encode the transmission ancestor in the extended clade as either a leaf label or a generic unknown sample, we lose information about the presence and size of blocks (i.e. chains of unsampled transmissions) on edges. The presence of transmission blocks can be inferred from the resulting extended topology, but their size is not explicitly encoded. Nevertheless, once a valid extended topology is obtained from the extended

CCD, details about blocks and their size can be reintroduced in a second step. Importantly, unlike decoupled summary approaches, the extended CCD first ensures a coherent joint structure before any further refinement and annotation is performed.

### Dataset Applications

To illustrate the practical differences between classical tree summary methods and our extended CCD framework, we analyse two empirical datasets featuring jointly inferred annotated phylogenies. The first dataset stems from a discrete trait phylogeographic analysis of the early spread of SARS-CoV-2 in Europe (Nadeau et al., 2021), based on a multi-type birth–death model (Scire et al., 2022). The second dataset concerns a measles outbreak in U.S. military bases following the withdrawal from Afghanistan in 2021 (Masters et al., 2023). While the original study employed a BEAST2 analysis, we reanalyse the data using the BREATH model (Colijn et al., 2026) in order to obtain posterior samples of transmission-embedded phylogenies. These two examples allow us to compare classical decoupled tree summaries with our extended CCD-based method in both phylogeographic and transmission settings.

### SARS-CoV-2 early transmission geography

The dataset of Nadeau et al. (2021) investigates the migration dynamics of SARS-CoV-2 prior to the spring 2020 border closures in Europe. The authors infer a partial transmission tree of the early pandemic while jointly reconstructing the geographic locations of ancestral lineages using a phylogeographic inference model. The discrete locations considered are Hubei (China), France, Germany, Italy, and “other European”. In addition to reporting a maximum clade credibility (MCC) summary tree, they analyse summary statistics describing inferred migration patterns among these locations. Here, we focus on the single-tree summaries and compare the published MCC tree with summaries obtained using the classical CCD, the HIPSTR approach as well as our extended CCD framework. We reuse the published posterior set of trees and refer to the original publication for more methodological details (Nadeau et al., 2021). For internal node states of the MCC, the regular CCD, and the HIPSTR trees, we use the most likely state as annotated by TreeAnnotator, corresponding to the state with highest marginal posterior probability at that node.

We begin by examining two metric multidimensional scaling (MDS) plots based on the classical RF metric and our extended RF metric. In Figure 10a), the five highlighted summary trees are distinct under both distance measures. Also shown is a sample of trees from the extended CCD, which clusters within the posterior point cloud in both plots, suggesting that the extended CCD faithfully represents the posterior structure of both topology and annotations. The extended RF distances appear to better capture the central tendency of all five summary trees. In contrast, the classical RF-based embedding results in these summary trees being more dispersed.

**Fig. 10:**
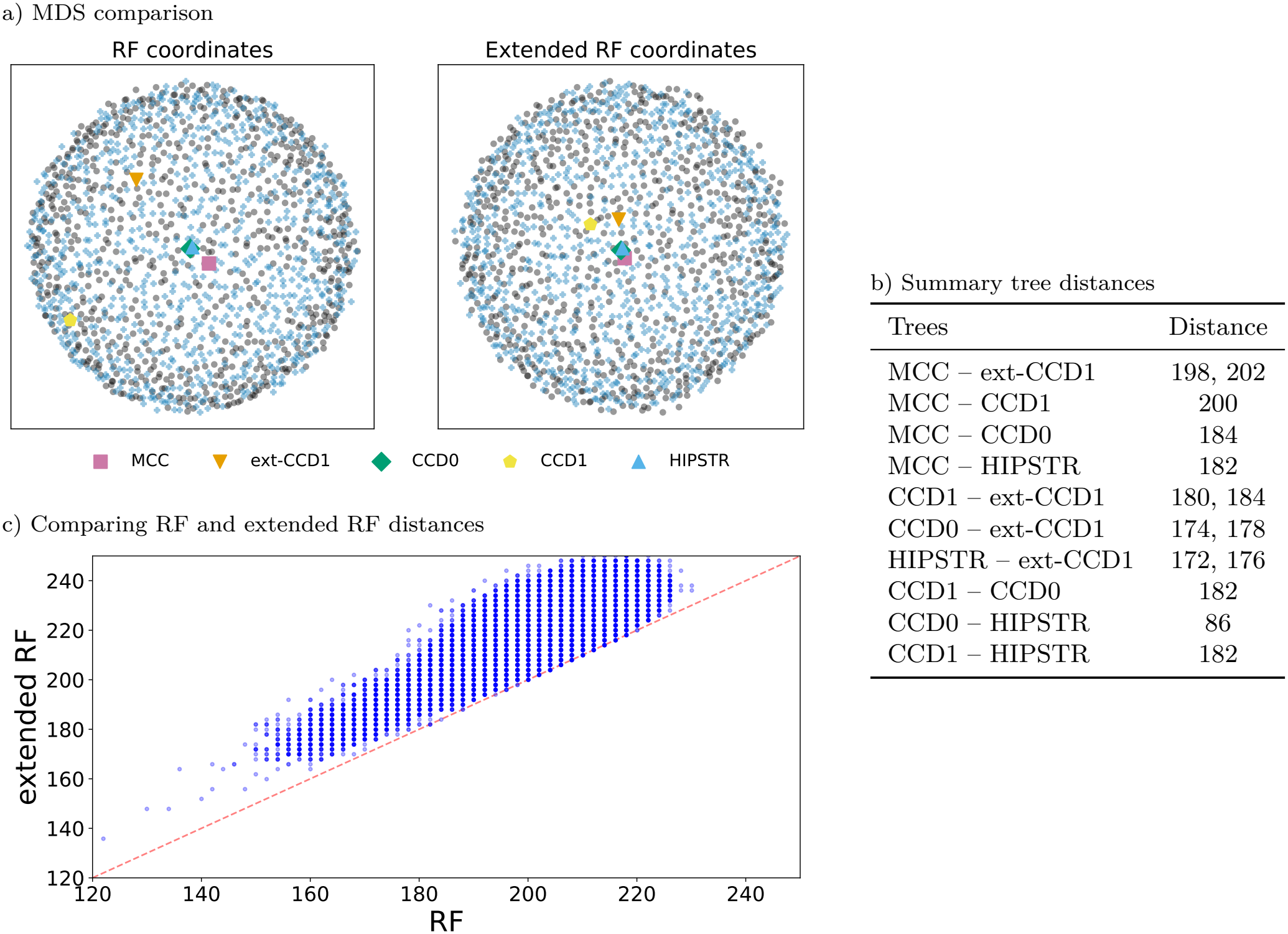
Summary tree comparison for the SARS-CoV-2 dataset. a) Metric MDS plots of 750 posterior trees (black circles) from the SARS-CoV-2 annotated trees, 750 trees sampled from the corresponding extended CCD (blue pluses), and the different summary tree options. We use the regular RF distance (left) as well as our extended variant (right). b) Pairwise distances between summary trees, two values indicate the RF and eRF distance if the two are different. c) Distribution comparison of the pairwise RF and eRF distances of the posterior tree sets that lead to the MDS plots in panel a).

In addition, we compare the pairwise distances between the five highlighted trees in Figure 10b). These comparisons reveal that the differences among the summary trees themselves arise mainly from differences in topology, that is, from their clade composition rather than from their geographic annotations. However, as illustrated by the MDS plot based on the extended RF metric, annotation differences substantially influence how the posterior trees are positioned relative to the respective summary trees. The extended RF metric captures joint topology-annotation structure, and this leads to its MDS plot placing the summary trees at the centre of the posterior sample.

The small pairwise distance between HIPSTR and CCD0-MAP in Figure 10b reflects that HIPSTR is equivalent to CCD0 without the expansion step described earlier.

### Comparing Summary Trees

We compare the extended CCD-MAP summary tree with the MCC tree published in Nadeau et al. (2021). Despite partial agreement, the extended CCD-MAP tree exhibits structural differences relative to the MCC tree. For example, the placement of the mostly German clade, the placement of the mostly Italian clade and also the splitting and different placement of the French clade differ between the two. These differences suggest that the extended clade structure captures aspects of the joint posterior that are missed when considering tree topology and annotations separately.

### Extended Majority Rule Consensus

In addition to the previously discussed summary trees, we also consider the majority rule consensus (MRC) tree, shown in Figure 12. This tree clearly illustrates the uncertainties present in the posterior distribution. Many of the differences observed between the MCC and extended CCD summary trees are reflected as unresolved clades (polytomies) in the extended MRC tree.

**Fig. 11:**
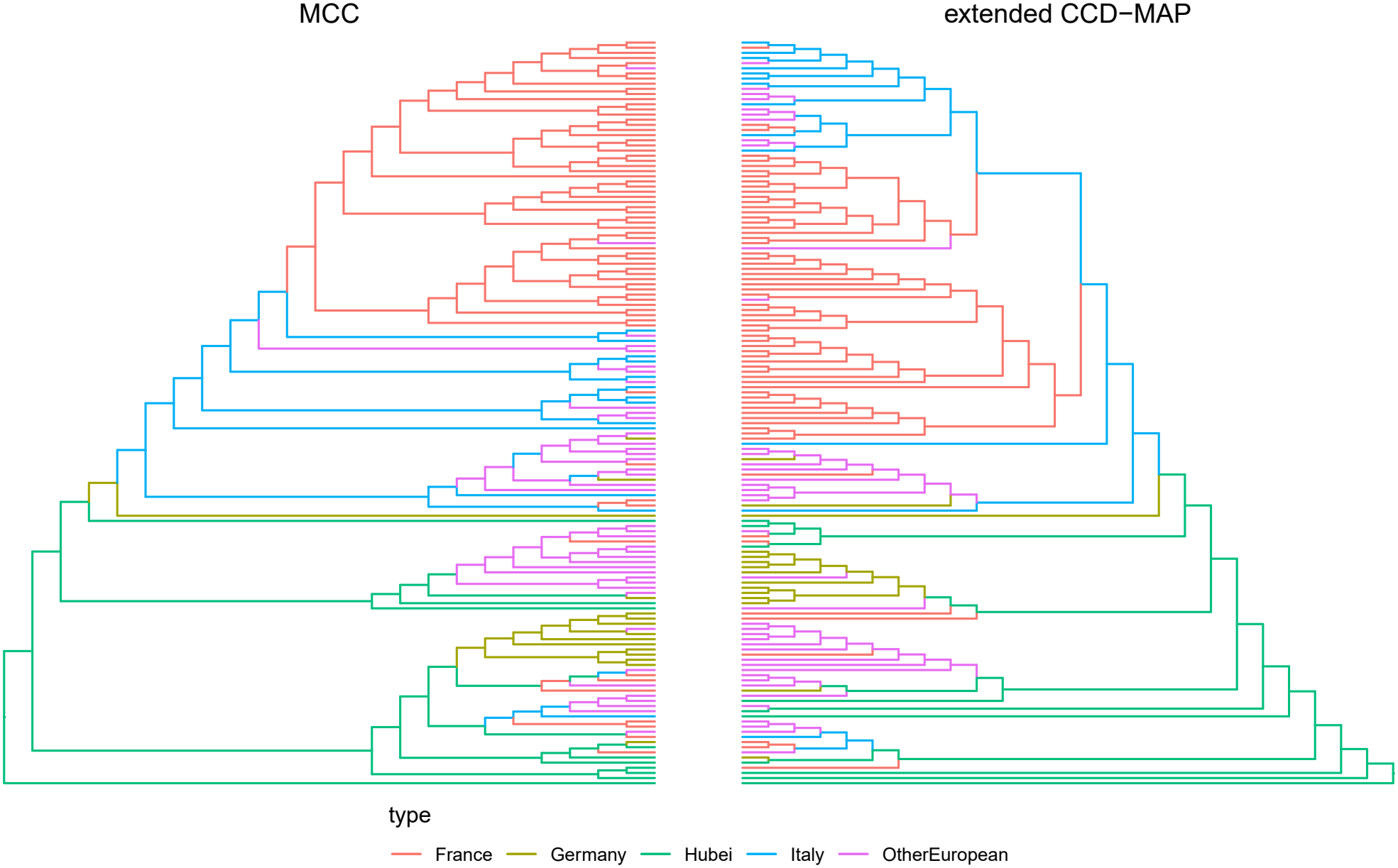
Comparing the MCC and the extended CCD-MAP tree topology without discarding any burn-in.

**Fig. 12:**
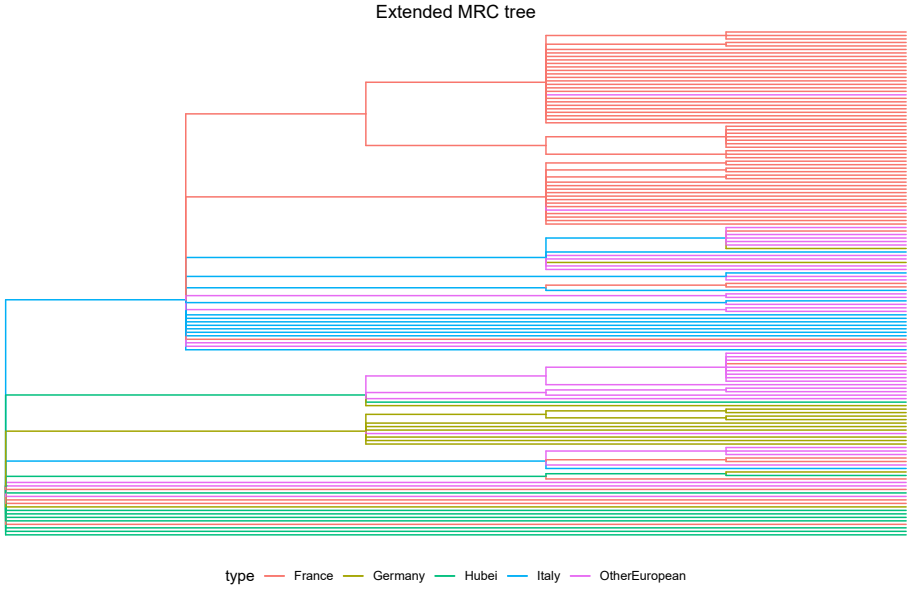
The extended MRC tree topology displaying the uncertainty of the full sample.

The MRC tree provides an intuitive visualization of posterior uncertainty, particularly when a fully resolved tree is not required. However, it is inadequate when a single, resolved topology with associated annotations is desired. In our example, substantial uncertainty remains in the posterior, and the choice of summary method determines which of the possible resolutions is highlighted.

### Measles outbreak

We analyse a measles outbreak that occurred among evacuees from Afghanistan during Operation Allies Welcome in 2021. This dataset was originally studied by Masters et al. (2023), where whole-genome data was used to reconstruct the transmission events. In this paper, we restrict our attention to the phylogenetic tree reconstruction. In the original analysis, the authors inferred a posterior sample of trees using BEAST 2.6.3 under a Bayesian skyline coalescent prior. We refer to the original manuscript and its supplementary material for full methodological details (Masters et al., 2023).

For our purposes, we do not aim to obtain an improved model fit under the BREATH model. Instead, we reuse the previously inferred parameter settings to conduct a computationally lighter BREATH analysis. Our objective is to examine the resulting posterior distribution of transmission trees rather than to optimize model specification. Consequently, we prioritize comparability and computational efficiency over a fully re-tuned model configuration. For this purpose, we use the subset of 41 individual sequences that were obtained in the original study.

### BREATH setup

We use a strict clock with rate 7.5 *×* 10^*−*4^ substitutions per site per year and we use the nucleotide substitution model *TIM* + *F* + *G*4 with the parameters provided in Masters et al. (2023). For BREATH specifics, we set the sampling fraction to be 0.97 and used the version of BREATH available on September 5, 2026 (v0.0.7). For the associated Gamma distribution we use a shape of 38 and a rate of 694, equivalent to a mean of 20 days and a standard deviation of *≈* 3.25 days. For the transmission hazard we use an expected infection count of 1.5. This reflects the assumption that there is considerable immunity to measles in adults, through either vaccination or previous infection. Infection timing is parameterized by a shape of 11.5 and a rate of 380, equivalent to a mean of 11 days and a standard deviation of *≈* 3.25 days. We fix the transmission population size to 0.05 which is equivalent to roughly 18 days.

We configure an MCMC analysis with a chain length of 200 000 000, using a thinning interval of 10 000. This results in a posterior sample of 20 001 trees. We visually inspected the parameter traces and ESS estimates and did not find any concerning trends. All of the files for this analysis are available online, see Data availability section.

### Who-infected-whom network

We begin by presenting the inferred who-infected-whom network in Figure 13. Such networks provide a more faithful representation of posterior uncertainty than any single summary tree and should generally be preferred when the aim is to summarize the full transmission posterior.

**Fig. 13:**
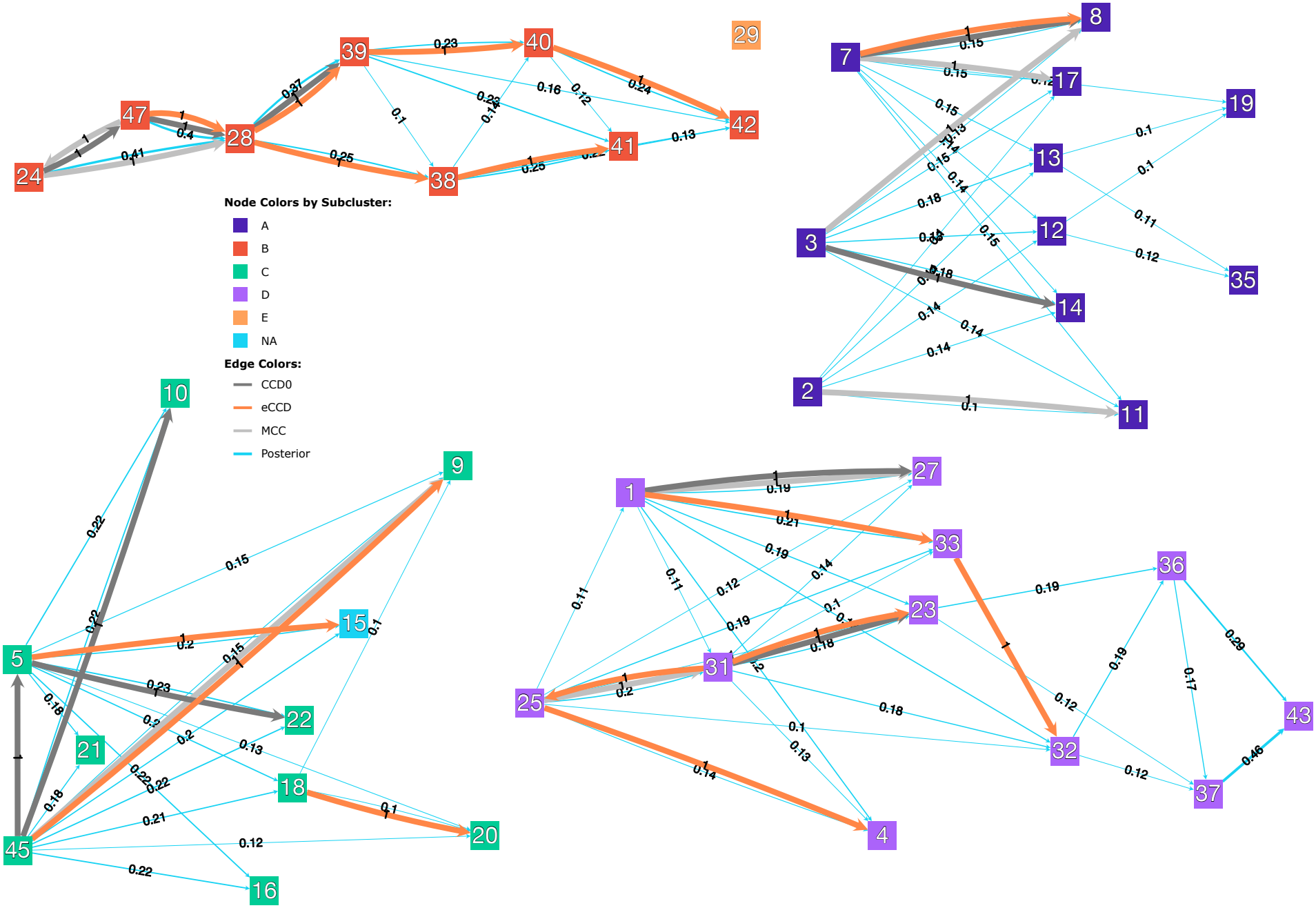
Inferred who-infected-whom network for the measles outbreak, including all direct transmission events with posterior support greater than 10%. Nodes are labelled by case number and coloured by previously inferred transmission sub-cluster (Masters et al., 2023). Edge colors correspond to the MCC (light grey), CCD0-MAP (dark grey), and extended CCD1-MAP (orange) summary trees and the posterior (light blue).

The who-infected-whom network recovers all five genetic clusters identified in the previous study, including one singleton (29) and one individual not assigned to any cluster (15). None of the summary trees violate this cluster structure by introducing edges between them. Overall, the two regular summary tree methods contain fewer direct transmissions (i.e. fewer edges) than our extended CCD1-MAP summary tree. For subcluster B (dark orange nodes) our method produces an almost complete transmission tree between the individuals only missing sample 24. For other subclusters, in particular subcluster A, the posterior edges express the uncertainty in these parts of the transmission scenario. A notable omission across all summary trees is the direct infection from case 37 to case 43, which has a very high posterior support of 0.43. In contrast, the MCC and CCD0-MAP summary trees introduce an edge between cases 24 and 47 despite its very low posterior support, and our extended CCD method introduces a transmission from case 31 to case 25 that was never sampled in the posterior. These variations highlight that any fully resolved summary tree necessarily represents only one particular realization of the underlying posterior transmission distribution, subject to the extent to which the chosen interpretation is able to faithfully capture its complex structure.

In Figure 14, we compare our extended CCD summary tree with an annotated CCD0 summary tree obtained by overlaying edge attributes onto a classical CCD topology using median summary statistics. Recall that the blockcount annotations are discrete and encode the transmission tree that is embedded into the phylogeny. Therefore, the median is not a suitable choice to summarize these values. We note that branch lengths for both summary trees can be overlayed separately using the default TreeAnnotator approach. This paper focuses on the extended topological structure, and we do not consider branch lengths further.

**Fig. 14:**
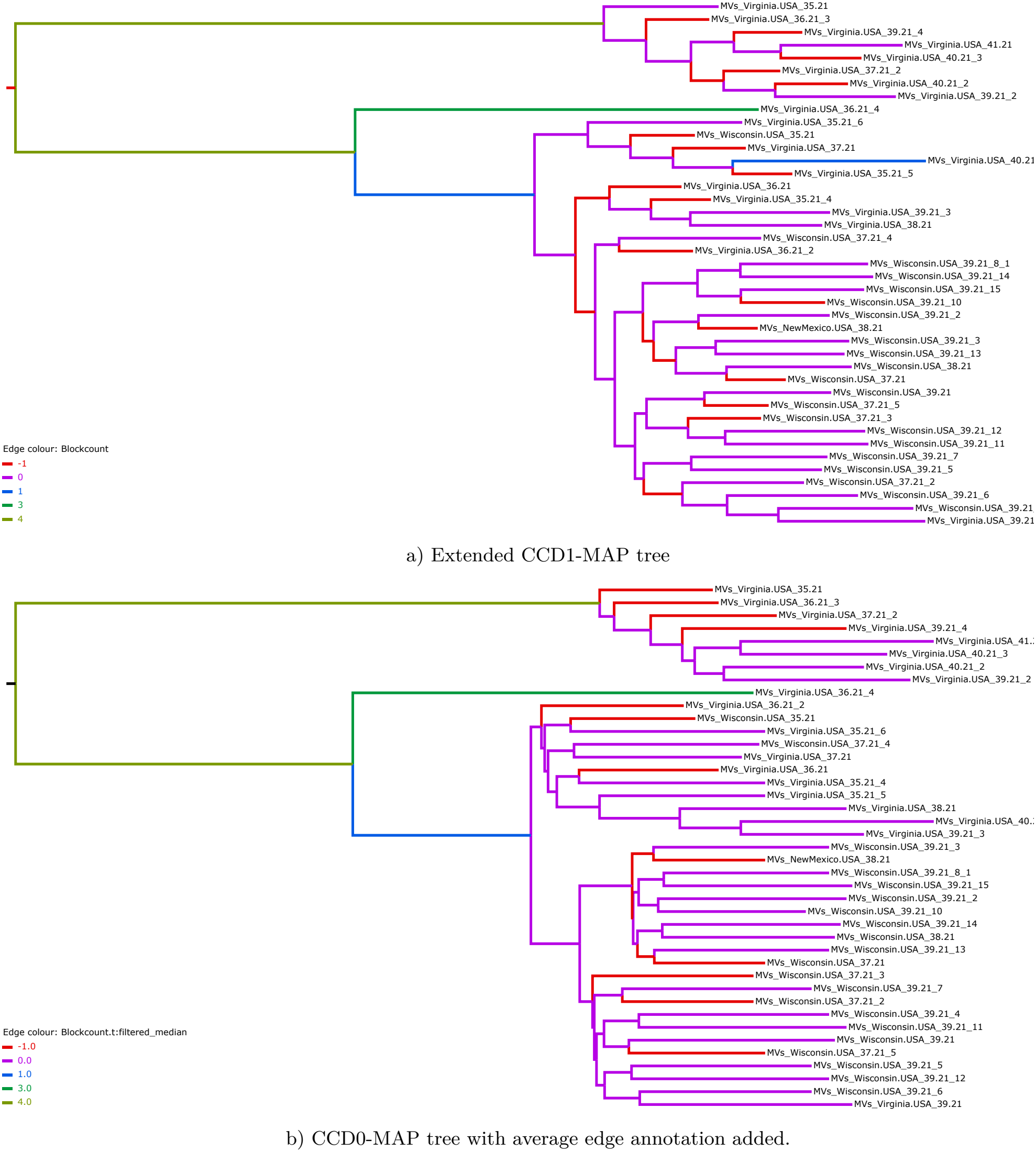
Comparing our extended CCD1-MAP tree to an annotated CCD0-MAP tree. Trees were visualized using IcyTree (Vaughan, 2017).

Both trees represent valid annotated trees under the BREATH model and we can observe several differences. For example, the smaller subtree below the root, corresponding to subcluster B in Figure 13, has a very different topology in the two trees. In the extended CCD1MAP tree we also observe a second long leaf branch with a block of size 1 on it.

In Figure 15 we provide two-dimensional MDS plots using the regular and extended RF metrics. We compare the posterior to a set of sampled trees from the extended CCD1 and the three previously presented summary trees. Under both metrics, the extended CCD1 samples and the posterior samples occupy similar and overlapping clouds of points. All three summary trees also lie within these clouds. However, under the extended RF metric, the extended CCD1 summary tree lies closer to the centre than under the regular RF metric.

**Fig. 15:**
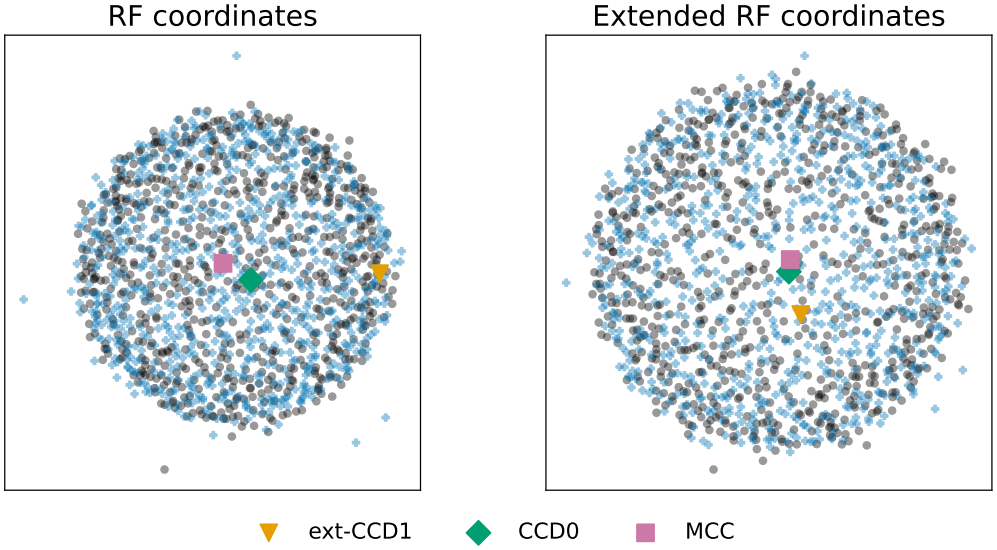
Metric MDS plots for the regular and the extended version of the RF metric. We compare a set of 750 posterior trees (black circles) from the BREATH analysis, 750 sampled trees from the corresponding extended CCD1 (blue pluses), the extended CCD1-MAP tree (orange triangle), the MCC tree (purple square), and the CCD0-MAP tree (green diamond).

Note that neither the regular CCD nor the MCC tree is guaranteed to be structurally valid under the BREATH model.

## Discussion

We introduced an extension of the classical clade that incorporates node or edge annotations that are inferred jointly with a phylogeny. This enables the extension of several classical clade-based methods: the conditional clade distribution, the Robinson-Foulds metric, and the majority rule consensus tree. The motivation for this development is the observation that, when topology and annotations are inferred jointly, decoupled summary procedures may yield representative trees that do not faithfully reflect the joint posterior, because the summary procedure treats topology and annotations separately even though they were inferred jointly. Proper sampling and marginalization from the joint posterior is, of course, always required for valid inference. However, a summary that preserves the joint dependence structure is specifically valuable when the question concerns the constraints and structure of that dependence itself.

The extended CCD framework ensures that extracted CCD-MAP trees correspond to valid annotated trees under the joint model. In the phylogeographic setting, we demonstrated that the classical two-step procedure can produce summary trees that violate model-implied constraints. The transmission tree setting highlights this distinction more sharply. Here, decoupled summaries may not merely reduce accuracy but produce structurally invalid annotated trees that correspond to no admissible transmission scenario. The extended CCD model is currently the only summary approach guaranteeing structural validity for these models.

Although the extraction of a MAP tree is a useful application, any single summary tree, including MCC, MRC, or CCD-MAP, necessarily condenses posterior uncertainty. This limitation is inherent to point estimates and not specific to our framework. Posterior support values displayed on a single representative tree may obscure uncertainty arising from alternative annotated topologies. The extended CCD representation addresses this by enabling analyses beyond a single tree. Extended clade probabilities can be computed directly, allowing assessment of support for annotated clades or partial tree structures. For example, in our extended settings with different root annotations, we can extract the extended CCD-MAP tree conditioned on each observed root annotation. In the transmission setting, we can extract and evaluate possible transmission configurations when conditioning on specific clade and infector pairs. Moreover, extended CCDs permit efficient sampling of valid annotated trees proportionally to their extended CCD probability. This can be useful for downstream analyses that require a collection of model-consistent trees rather than a single representative.

The extended clade framework naturally induces an extended RF tree metric. This metric retains the settheoretic interpretation and computational tractability of the classical RF metric while incorporating annotations. This complements previous tree-edit-based generalizations of the RF metric (Briand et al., 2020) and opens avenues for the study of annotated tree metrics. Furthermore, such a metric is a valuable tool to analyse posterior or bootstrap collections of hundreds of trees through means such as MDS plots or distance distributions. We can also use the extended clade framework to extend tree summary strategies such as the strict and majority-rule consensus trees. These trees are particularly useful for representing posterior uncertainty as an only partially resolved consensus. The extended clade framework could similarly be used to extend the existing MCC and HIPSTR (Baele et al., 2025a) regular clade based tree summary frameworks.

Beyond the two motivating applications, the extended clade framework is readily generalizable to other classes of structured trees for which principled summary methods remain underdeveloped. Potential applications include sampled ancestor trees under fossilized birthdeath models (Gavryushkina et al., 2014), stratigraphic range trees (Stolz et al., 2025), and possibly structured network-like objects arising in horizontal gene transfer or language evolution, for which inference methods already exist (Müller et al., 2020; Neureiter et al., 2022).

We now discuss several limitations of the current framework. First, tree topology and branch lengths are rarely treated jointly: theory and tooling tend to focus on one or the other. Continuous branch lengths are naturally handled in tree spaces such as BHV (Billera et al., 2001) or *τ*-space (Gavryushkin and Drummond, 2016), while discretised branching times are handled in tree spaces such as RNNI (Gavryushkin et al., 2018) or RSPR (Collienne et al., 2024). However, none of these have been widely adopted in standard phylogenetics software as tree summary tools. Extending our or other frameworks to summarise branch lengths and topology jointly, rather than sequentially, is therefore a promising but substantial direction for future work, warranting its own dedicated study. Second, the CCD0 expansion step is model-specific: compatibility constraints between extended clades must be defined for each annotation type, and no general solution for clade and clade split compatibility exists for arbitrary models. Third, incorporating annotations increases the total number of possible clades. Each classical clade now appears in up to |*A*| annotated variants. Consequently, larger posterior samples are required to achieve reliable extended clade and extended clade split frequency estimates. Fourth, in the transmission tree setting, information about the size and placement of unsampled transmission chains (blocks) is not retained in our extended clade representation, as discussed in the “Transmission Tree Reconstruction” subsection. While this information can be reintroduced in a subsequent step, it is not captured by the extended CCD itself. Finally, the current implementation is limited to two inference frameworks: discrete traits, as in phylogeography, and transmission-embedded phylogenies as inferred under the BREATH package for BEAST 2. Extending the framework to other tools and annotation types is in principle straightforward but requires careful handling of non-standardized tree formats and annotation encodings, which vary considerably across inference software.

More broadly, our framework connects to ongoing work on posterior entropy and dissonance measures (Lewis et al., 2016; Berling et al., 2025), rogue taxon identification (Klawitter et al., 2024), and confidence interval estimation (Klawitter and Drummond, 2026) within classical clade-based CCDs. With our extended clade framework it is in principle straight-forward to adapt these for annotated trees, representing a natural avenue for future work. Furthermore, our framework can be combined with the work presented in (Takazawa et al., 2026) to get a more refined version of the extended MRC summary tree. Along these lines, a refinement of our extended RF metric is also a natural point for future work, for example using the approaches from (Smith, 2021). As joint inference models of annotated trees or networks become more prevalent, the extended clade framework provides a principled and extensible foundation for developing the corresponding summary and analysis tools.

## Acknowledgements

We would like to thank Olivier Gascuel for helpful comments and discussions during early stages of this project. We thank Tanja Stadler, Tim Vaughan and the rest of the Computational Evolution group in Basel for being a critical audience and providing valuable feedback to early versions of the results. We would like to thank Alexei Drummond for helpful comments and suggestions. We are grateful to the Associate Editor, Jordan Douglas, and the two reviewers, Luiz Max Carvalho and an anonymous reviewer, for their constructive comments and suggestions, which helped improve the paper. This work was funded by the Federal Government of Canada’s Canada 150 Research Chair program.

## Data availability

The two example datasets are freely available online with their respective publications (Nadeau et al., 2021; Masters et al., 2023). We have also deposited our analysis scripts and the required original data at https://doi.org/10.5281/zenodo.22663643. Implementation of the extended CCD methods can be found at https://github.com/Lars-B/brokilon. The who-infected-whom network Figure 13 was generated with the app available at https://github.com/Lars-B/contagion.

